# Learning ‘*The Knowledge’*: How London Taxi Drivers Build Their Cognitive Map of London

**DOI:** 10.1101/2021.06.04.447168

**Authors:** Eva-Maria Griesbauer, Ed Manley, Jan M. Wiener, Hugo J. Spiers

## Abstract

Licenced London taxi drivers have been found to show changes in the grey matter density of their hippocampus over the course of training and decades of navigation in London (UK). This has been linked to their learning and using of the ‘*Knowledge of London*’, the names and layout of over 26,000 streets and thousands of points of interest in London. Here we examined the process of how this knowledge is acquired and we detail key steps that include: systematic study of maps, travel on selected overlapping routes, the mental visualisation of places and the optimal use of subgoals. We provide the first map of the street network covered by the routes used to learn, allowing insight into where gaps in the network exist. The methods could be widely applied to aid spatial learning in the general population and may provide insights for artificial intelligence (AI) systems to efficiently learn new environments.

## Introduction

The ability to navigate an environment depends on the knowledge of that environment. This knowledge can be gained in multiple ways, such as via instructions on GPS devices, memorising a cartographic map, or through exploration. The knowledge formed can vary from very imprecise to extremely accurate, depending on the complexity of the environment, the level of exposure to the environment and individual differences (Schinazi et al, 2013; Weisber et al., 2014; Weisberg and Newcombe, 2016; Ekstrom et al., 2018). Over the last decades, there has been increasing interest in understanding how different methods for learning impact the acquisition of spatial knowledge (e.g. Dahmani & Bohbot, 2020, Hejtmánek, et al., 2018; Gardony et al., 2013; Münzer et al., 2012; Ishikawa, Fujiwara, et al., 2008; Münzer et al., 2006; Siegel, & White, 1975; Streeter & Vitello, 1986, Balaguer et a., 2016) and how individuals differ in their capacity to learn to navigate new environments (Weisberg and Newcombe, 2016; Coutrot et al., 2018; Coutrot et al., 2019; Coutrot et al., 2020; Newcombe, 2018; Burles and Iaria, 2020).

Despite GPS devices being a preferred method of navigation for many (McKinlay, 2016), the increased use of GPS devices appears to have a negative impact on spatial memory (Dahmani & Bohbot, 2020; Ruginski et al., 2019) and is associated with habitual learning of a particular route (Münzer et al., 2006). In contrast to GPS-based instruction-guided navigation, ‘map-based navigation’ (relying on memory for the map) has been found to support spatial learning, knowledge acquisition of the environment and improved flexible navigation performance (e.g. Ishikawa, Fujiwara, et al., 2008; Münzer et al., 2012; Münzer et al., 2006). Such flexible navigation relying on long-term memory is associated with the construction of a cognitive map, which stores the allocentric information about the structure of the environment enabling shortcuts and efficient detours around unexpected obstacles (Tolman, 1948; O’Keefe and Nadel, 1978).

A range of evidence indicates that within the brain the hippocampus provides a cognitive map of the environment to support memory and navigation (O’Keefe & Nadel, 1978; Gahnstrom & Spiers, 2020; Epstein et al., 2017) and damage to the hippocampus disrupts navigation (Morris et al., 1984; Spiers et al., 2001). Hippocampal neurons encode spatial information (O’Keefe and Nadel, 1978) and for a selected group of individuals, who spend their daily lives navigating using map-based recall of space, their posterior hippocampal grey matter volume increases with years of experience and is larger than in the general population (Maguire et al., 2000). These individuals are licensed London taxi drivers. Here, we explore how they gain the vast knowledge required to navigate London, which appears to drive the changes in their hippocampus (Woollett and Maguire, 2011).

Licensed London taxi drivers are unusual among taxi drivers. They are able to mentally plan routes across an environment that contains more than 26,000 streets within the six-mile area around Charing Cross, the geographic centre of London (A to Z from Collins The Knowledge, 2020). They are required to have sufficient knowledge to also navigate main artery roads in the suburbs. This area covers almost 60,000 roads within the circular M25 (The London Taxi Experience - The Knowledge, 2020; numbers may vary depending on sources, road types and the definition of the boundary of London). What makes licenced London taxi drivers unique is that they have to accomplish this using their own memory, without relying on physical maps or navigation aids. They are also the only taxi drivers allowed to pick up customers when hailed in the street, due to their license to operate. In the rest of this article, we refer to them as London taxi drivers, but readers should note that our analysis pertains only to licensed taxi drivers.

Following the discovery of changes in hippocampal size in London taxi drivers by Maguire and colleagues (2000) numerous studies have explored their brain function and cognition. Neuroimaging have provided further evidence of structural changes in their hippocampus (Maguire et al., 2006; Woollett, Spiers & Magure 2009; Woollett & Maguire, 2011). London taxi drives show a greater posterior hippocampal grey matter volume compared to London bus drivers, who also navigate London daily, but take the same routes (Maguire, Nannery & Spiers, 2006). Examining brain changes longitudinally, Woollett and Maguire (2011) found that an increase in the posterior hippocampus after the years spent training to learn what is known as the Knowledge of London and pass the exam required to become a licensed taxi driver (Woollett & Maguire, 2011). Notably, those who failed to qualify did not show a change in their hippocampal size. Acquiring the Knowledge of London seems to come at a cost of learning and retaining new visuo-spatial information, which co-occurs with a concurrent volume decrease in the anterior hippocampus (Maguire, Woollett & Spiers, 2006; Woollett & Maguire, 2009; Woollett & Maguire, 2012). Functional neuroimaging studies have shown engagement of their posterior hippocampus when verbally recalling routes (Maguire et al., 1997) and at the start of the route when navigating a highly detailed virtual simulation of London (Spiers & Maguire, 2006; Spiers & Maguire, 2007). Other research with London taxi drivers has revealed insight into spontaneous mentalizing (Spiers and Maguire 2006b), emotions during navigation (Spiers and Maguire 2008), the neural basis of driving a vehicle (Spiers and Maguire, 2007b) and the route planning process (Spiers and Maguire, 2008). London taxi drivers have also been shown to be better than non-taxi drivers at learning new routes (Woollett and Maguire, 2009).

Despite the numerous studies exploring London taxi drivers, little attention has been paid to how London taxi drivers learn and memorize the layout and landmarks in London (Skok, 1999). Many questions arise when considering this. How is their exploration structured? What do they study when examining maps. How are map and physical travel experience integrated? What role does mental imagery play in aiding their learning? How do they exploit the hierarchical structure of London’s layout? Are major roads mastered before minor roads? In this observational report we provide the first investigation of London taxi driver’s learning process and the methods and techniques that enable them to retain and use such a large amount of real-world spatial information for efficient navigation.

## Methods

To understand the learning process of taxi drivers, different types of sources of information have been consulted. These sources included (a) a semi-structured interview (ethics approval was obtained under the ethics number CPB/2013/150) with a teacher from a London *Knowledge school* (here referred to as K.T. for ‘*Knowledge Teacher*’), (b) an email exchange with Robert Lordan, the author of ‘The Knowledge: Train Your Brain Like A London Cabbie’ (Lordan, 2018), (c) an open introductory class of the *Knowledge of London* and regular scheduled classes for current students, (d) school specific study material and (e) online information from the TfL (Learn the Knowledge of London, Transport for London, n.d.; Electronic blue book, 2019).

The interview with the teacher from the *Knowledge school* was audio-recorded and transcribed. The transcription of the interview can be found in **Appendix A**. The teacher gave written consent for the content of this interview to be cited and published. Additionally, attendances of *Knowledge school* training classes, including an introductory class and several classes with more advanced students, allowed to observe and understand the training process in more detail.

The information collected from these sources was systematically reviewed to report on (a) the ways spatial information is structured and presented for the learning process, (b) the techniques and methods used to learn this spatial information and (c) the ways how this information is tested and the later perception of this knowledge as a taxi driver. A summary for each of these categories was created, starting with verbal reports (interview (**Appendix A**), *Knowledge school* classes). This information was cross-referenced with and extended by unreported information from other, published or official sources (e.g. study material, online booklets by TfL). Additionally, where verbally reported and based on official sources, visuals of essential places of interest and routes were created.

## Observations

Taxi drivers in London have to demonstrate a thorough Knowledge of London within the six-mile radius originating at Charing Cross (see Figure 1a) to earn the green badge that qualifies them to drive a ‘black cab’ taxi (Electronic blue book, 2019). Within this area, taxi drivers are expected to plan a route (i.e. the ‘runs’) based on the shortest distance between any two potential places of interest (i.e. the ‘points’) their customers might travel from or to, such as restaurants, theatres, hospitals, sports centres, schools or parks (cf. Electronic blue book, 2019, for a complete list). Taxi drivers are also expected to name all roads or streets that are part of that run in the correct, sequential order, including travelling instructions, such as turns (Electronic blue book, 2019).

**Figure 1.**
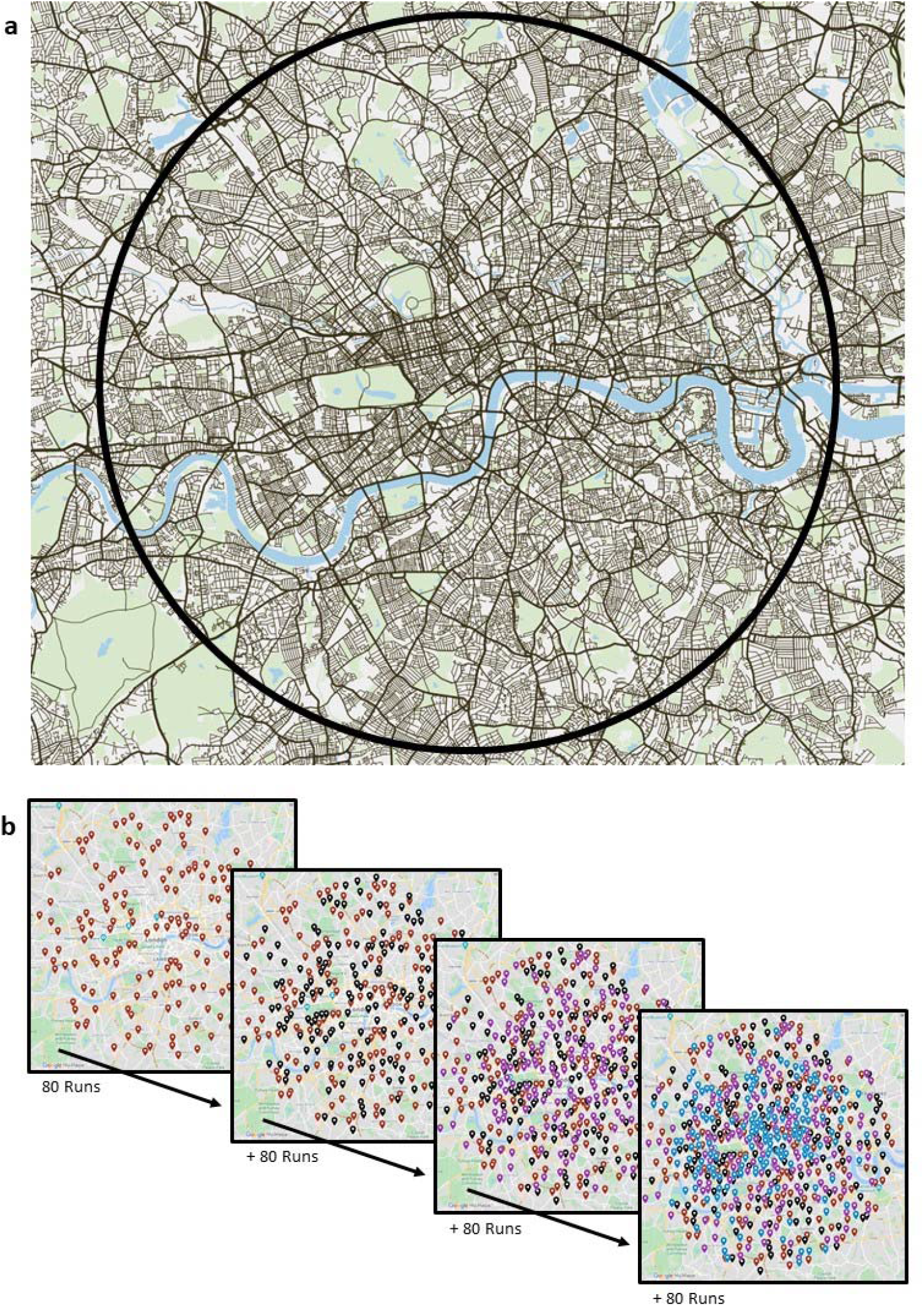
The Knowledge of London and the Blue Book. (a)London taxi driver students are expected to learn the street network and all potential points of interest within the six-mile radius around Charing Cross (black circle), which is called the ‘*Knowledge of London*’. (b) To support the learning process of this area, the Blue Book has been put into place. It contains 320 origin-destination pairs and the shortest route (i.e. ‘*run*’) connecting those pairs. When mapped chronologically in groups of 80 *runs*, the network of origin-destination pairs starts overlapping and becomes denser. **Red**: the first layer of the first 80 origin-destination Paris. **Black**: The second layer of the origin-destination pairs for *Runs* 81-160. **Purple**: The third layer of origin-destination pairs for *Runs* 161-240. **Blue**: The final layer of the last 80 origin-destination pairs for *Runs* 241-320. Map sources: (a) <u>Mapbox</u>, (b) <u>My Maps</u> by Google Maps.

Historically, the exact roots of the *Knowledge of London* are unclear as written evidence is mostly missing. The first licenses and regulations for horse-driven carriages date back to the early 1600s by Oliver Cromwell (London Metropolitan Archives, 2013; Newton, 1857; June 1654: An Ordinance for the Regulation of Hackney-Coachmen in London and the places adjacent, 1911; Lordan, 2018). However, in 1851 the Great Exhibition in Hyde Park revealed incompetent navigation skills of the carriage drivers of those days. These initiated a series of complaints and forced authorities in the following years to set up stricter qualification requirements for drivers to test their knowledge of important streets, squares and public buildings (Lordan, 2018; A to Z from Collins - The Knowledge, 2020; The Knowledge, London’s Legendary Taxi-Driver Test Puts Up a Fight in the Age of GPS, 2014). This scheme was officially introduced in 1865 (Learn the Knowledge of London, Transport for London, n.d.). The requirements in relation to the content of the *Knowledge* have since hardly changed and remained in place (The Knowledge, 2020) despite the technological innovations that have produced navigation aids, such as GPS devices, that facilitate and guide navigation. The following sections will outline how this is achieved by taxi drivers.

### Presentation of Spatial information in Knowledge Schools

To help students to acquire the fundamentals of the *Knowledge of London*, the Blue Book (the origin of this name is unclear) was designed, which, in its current form, was put into place in 2000 (interview with K.T., **Appendix A**). It contains 320 origin-destination pairs, their corresponding *runs*, as well as additional *points* related to tourism, leisure, sports, housing, health, education and administration (Electronic blue book, 2019). In total, there are about 26,000 different streets and roads (Eleanor Cross Knowledge School, 2017) and more than 5,000 *points* (Full set of Blue Book Runs, 2020) listed in the *Knowledge* schools’ versions of the Blue Book. However, this knowledge is incomplete. By the time students qualify, they will have extended their knowledge to identify more than 100,000 *points* (The London Taxi Experience - The Knowledge, 2020) in a street network of about 53,000 streets (OS MasterMap Integrated Transport Network, 2018). This covers not only the six-mile area, but extends to all London boroughs, including major routes in the suburbs.

The 320 origin-destination pairs of the Blue Book with their corresponding *runs* are structured into 20 lists of 16 pairs each, which are designed to systematically cover the six-mile radius: In a chronological order, as listed in the Blue Book, the majority of origin-destination pairs have an origin in the same postal districts as the destination of the previous origin-destination pair and spread across London throughout each list (Electronic blue book, 2019). When mapped in layers of four, the first 80 *runs* (i.e. five lists) provide an initial rough coverage of London. This coverage becomes denser with each of the remaining three layers that are shifted slightly against each other to fill in the gaps (**Figure 1b**).

Each of the origins and destinations in the Blue Book also require students to learn the nearby environment within the quarter mile range. That area around a Blue Book *point* is called the ‘*quarter mile radius*’, or in short: the ‘*quarter-miles*’ and is considered as ideal for learning small areas of the environment without overloading students with information (interview with K.T., **Appendix A**; Learn the Knowledge of London, Transport for London, n.d.; Electronic blue book, 2019). For the first and most famous *run*, which connects Manor House Station to Gibson Square, the quarter-mile radius is illustrated in **Figure 3a**. It contains about 8 additional *points*, numbered 1-8. These are chosen by each *Knowledge school* individually and can differ between schools. The additional *points* serve as initial motivation for students to explore the quarter-miles and learn which streets link these points to each other. Knowledge of the remaining, unmentioned *points* in the area will be obtained by each student gradually as they progress through the *Knowledge of London* by studying maps and exploring the quarter-miles in person.

**Figure 2.**
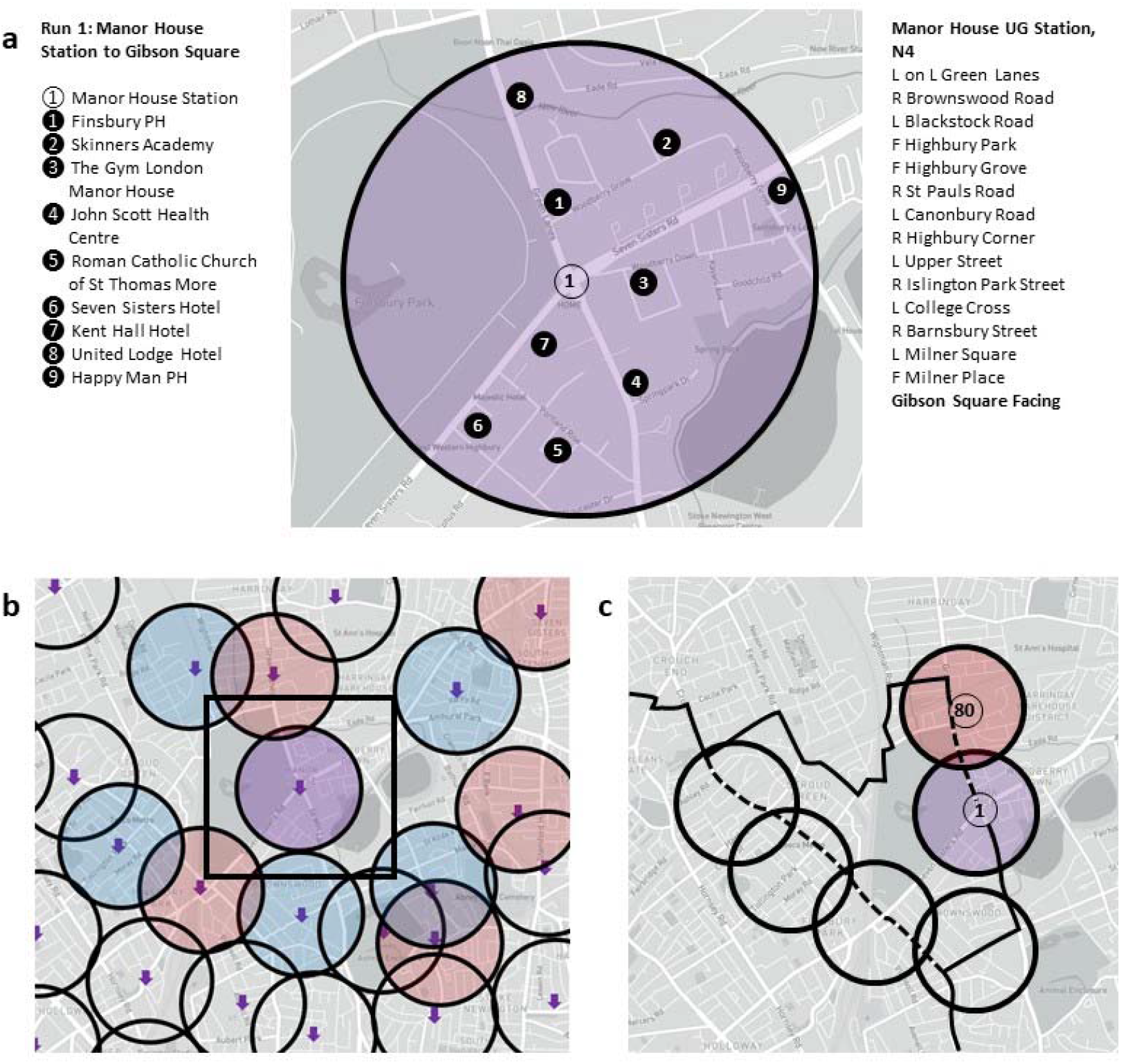
The points of the Blue Book. Each origin-destination pair of the Blue Book is presented in relation to its quarter-mile area. The origin of a *run*, here *Run* 1 (a), Manor House Station, and the corresponding quarter mile radius (black circle) with additional 8 other points of interest (numbered 1-8) are marked in a map. Labels are provided in a legend (left) and the most direct route (i.e. ‘*run*’) to the destination, including driving instructions (L on L: leave on left, L: left, R: right; F: forward) are listed on the right. The dense network of origin-destination pairs (b) results in an overlay of the neighbouring quarter-mile ridii (black circles around purple arrows). For Manor House Station (purple circle) neighbouring quarter-mile origins and destinations are highlighted in blue and red, respectively. These quarter-miles are covering the six-mile radius in London by linking places of interest through linking *runs* (c) as indicated by the dashed lines connecting *Run* 1 from Manor House Station and *Run* 80, ending at Harringay Green Lanes Station. Source: Figures are based on learning material from <u>Taxi Trade Promotions</u>.

**Figure 3.**
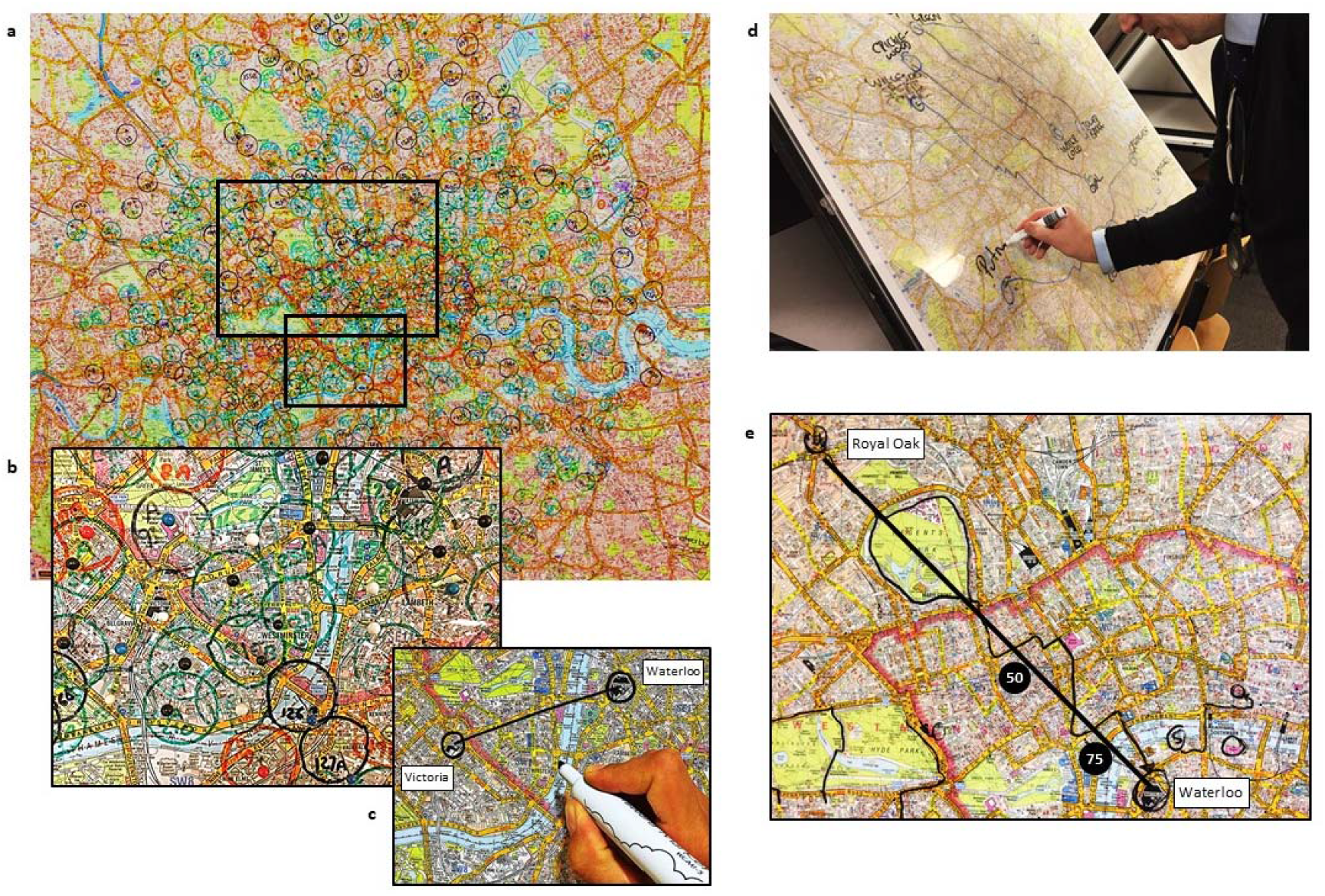
Example of Knowledge school material in use. In *Knowledge schools*, wallpaper maps (a) are used to illustrate the coverage of London within the six-mile area by the quarter-mile radii (b). These maps support the learning of relations between two places and clear up misconceptions such as Victoria being located further north than Waterloo, which is owed to a change in direction of the River Thames (c). ‘The cottoning up of two points’, a piece of string that is used to create a direct line between the points, is a common method to help with directional studies (c) and planning the most direct routes (d, e). Additionally, students use 50% and 75% markers along the direct line (e) to create subgoals that help to plan the *runs*. Source: Knowledge Point School, Brewery Road, London, UK.

Mapping the origin-destination pairs with their corresponding quarter-miles, highlights how the areas locally link to each other (**Figure 3b**). To create such an overlap that sufficiently covers the whole six-mile area around Charing Cross (also see **Figure 3a**), 640 *points* are required, thus explaining the total number of 320 Blue Book *runs*. Since each *point* is closely surrounded by nearby origins and destinations of other *runs*, information is provided about how an area can be approached from or left in different directions. For Manor House (**Figure 3b**) these *points* have been indicated by blue and red quarter-miles for nearby origins and destinations, respectively, in **Figure 3b**. To visualise this information across the entire six-mile area of London and keep track of their progress whilst learning the Blue Book, trainee taxi drivers mark the origins and destinations, including the quarter-miles, in a large, all London map (**Figure 3a** and **b**, source: Knowledge Point Central, Brewery Road, London, UK).

Studying maps by visualising the topological relationship between areas also helps to avoid misconceptions about the city’s geography that could lead to mistakes in route planning. For instance, deviations from the more generally perceived west-east alignment of the river Thames can cause distortions (cf. Stevens & Coupe, 1978). Often Victoria station, located north of the river, is incorrectly perceived further north than Waterloo Station, which is on the southern side of the river, but further east then Victoria (see **Figure 3c**). This misconception is due to a bend of the river Thames, that causes the river to flow north (instead of east) between Victoria and Waterloo.

In the Blue Book, the 320 *runs* connect the origin-destination pairs through the route along the shortest distance for each pair (Electronic blue book, 2019). These pairs were chosen to create *runs* that are about two to three miles long and mainly follow trunk or primary roads. Here, trunk roads are the most important roads in London after motorways, providing an important link to major cities and other places of importance, with segregated lanes in opposite directions (Key:highway, 2020). Primary roads are defined as the most important roads in London after trunk roads, usually with two lanes and no separation between directions, linking larger towns or areas (Key:highway, 2020). Since these are often printed in orange and yellow in paper maps, taxi drivers also refer to them as ‘*Oranges and Lemons*’ (interview with K.T., **Appendix A**). Trainee taxi drivers visualise these *runs* on all London maps to learn and practice recalling them (**Figure 3d**, credit: Knowledge Point Central, Brewery Road, London, UK). *Knowledge schools* provide the 320 *runs* for the *points* of the Blue Book but encourage students to plan these *runs* before checking the up-to-date solution. To plan a *run* using the shortest distance and avoid major deviations (as required for the examinations), drawing the direct line (i.e. ‘*as the crow would fly*’) or spanning a piece of cotton between the *points* is essential (**Figure 3e**). This so-called ‘*cottoning up*’ also helps students to learn relations between places (**Figure 3c**) and visualise the map to find ways around obstacles, such as Regent’s Parks, or to select bridges for crossing the river **(Figure 3e**) during the ‘*call out*’ of the *run* (i.e. the recall of the street names in order along shortest route without using a map). Additionally, it provides opportunities to set subgoals, the ‘*50% and 75% markers’*. These markers are set where the line coincides with major roads or bridges, about halfway or three quarters along the line. These distances are guidelines only, and sometimes bullets are set at other distances for streets and places along the direct line that facilitate planning in stages. These markers help students to stay close to the direct line, whilst breaking down longer *runs* in smaller sections and reduce the number of steps they have to plan for at a time **(Figure 3e**). Due to one-way streets and turning restrictions, *reverse runs* from the initial destination to the initial origin can differ. Therefore, the streets and roads cannot simply be called in reverse order but have to be learned separately (**Figure 4**).

**Figure 4.**
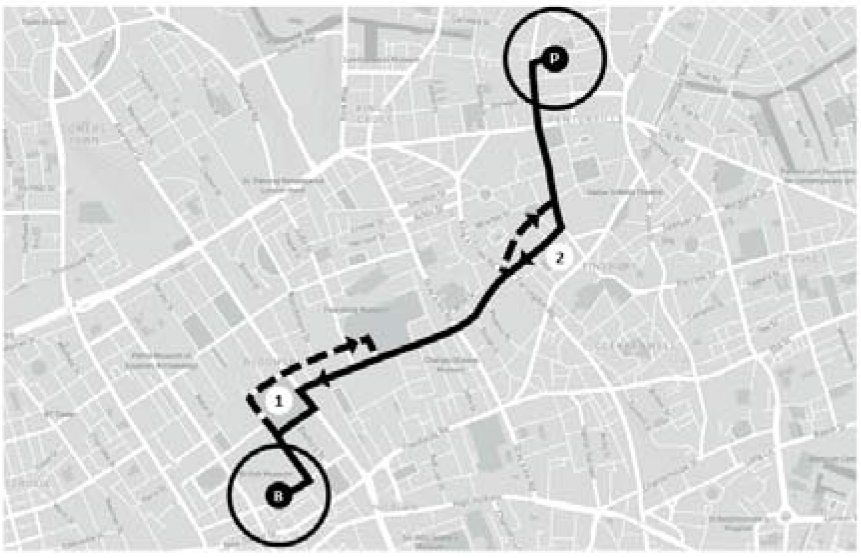
Runs and reverses runs. Due to one-way systems or turning restrictions, some *runs* differ when planned in reverse (dashed line), not allowing to simply invert the original sequence of streets taken (black line). This is the case for the *run* from Islington Police station (P) to the British Museum (B). When reversed, the one-way systems at Russell Square (1) and at Margery Street (2) require adaptation to traffic rules, resulting in differences between the *runs* and its reverse *run*. Figure is based on learning material from <u>Taxi Trade Promotions</u>.

The *runs* of the Blue Book form a network of routes that covers the six-mile area centred around Charing Cross (**Figure 5a**). However, the coverage of the London street network by the Blue Book *runs* systematically varies in density with respect to the distribution of *points* and the complexity of the street network: At its boundaries **(Figure 5b**) this network is less dense than in central London, where the *runs* are also overlapping more often (**Figure 5c**). This also reflects that more *points* are located closer to the centre of London, whereas residential areas are more likely to cover larger regions at the boundaries of the six-mile radius. Similarly, areas of London with a more regular street network, such as in Marylebone and Fitzrovia, are covered by less *runs* (**Figure 5d**) than areas with a more complex and irregular street network, such as South Kensington and Chelsea (**Figure 5e**). These might require more practice to learn.

**Figure 5.**
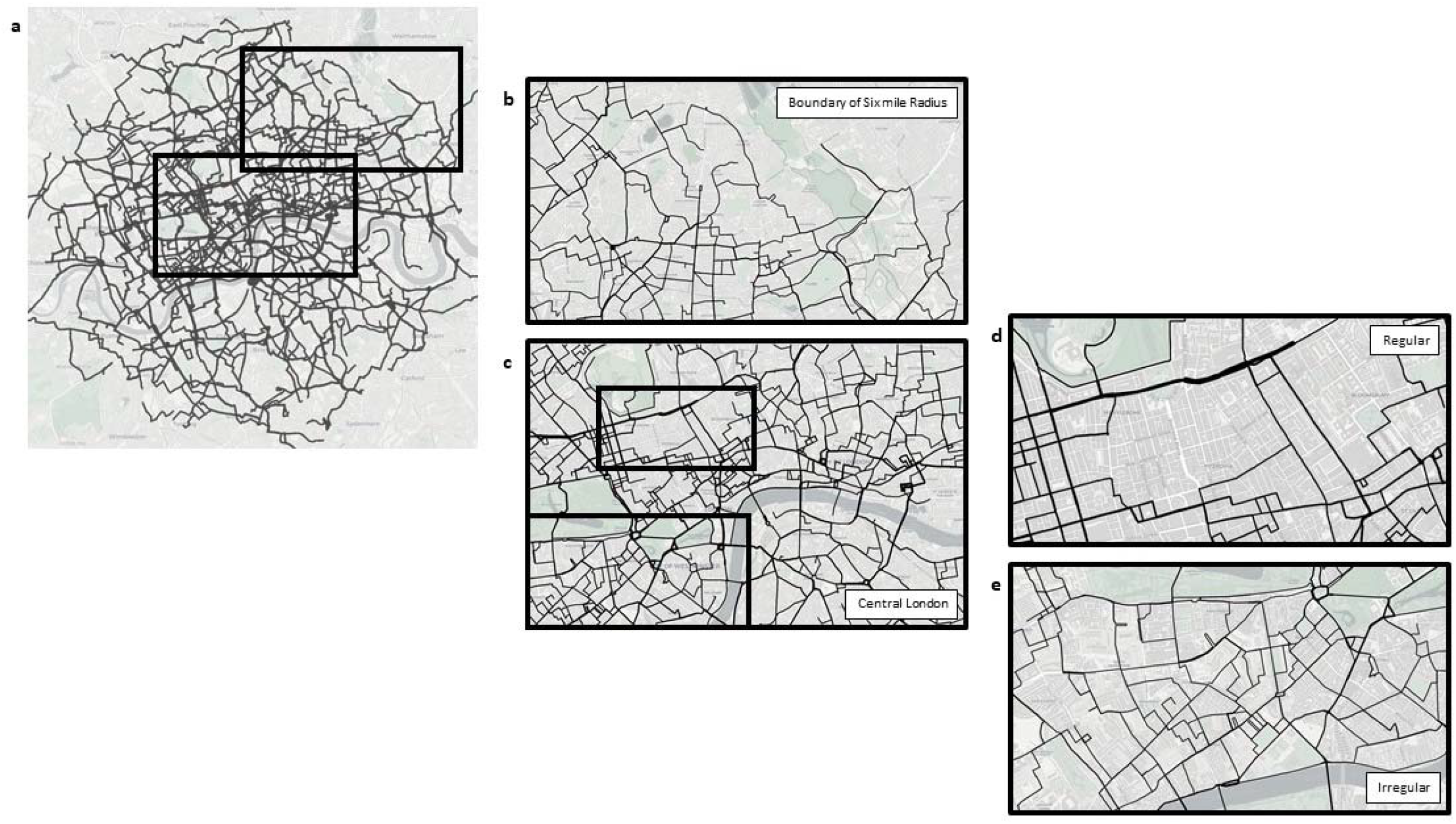
Network of Blue Book runs. A visualisation of the 320 *runs* that connect the corresponding origin-destination pairs of the Blue Book forms a dense network of routes that overlaps, similar to the quarter-mile radii (a). Across the network, density varies and is less dense closer to the six-mile boundary (b) then in central London (c). This overlap also shows that more routes *run* through areas with higher irregularity in the street network (d) than areas of a more regular street network (e) in central London. Source: Adapted from Blue Book mapping by Prof Ed Manley, University of Leeds

The Blue Book *runs* focus on connecting origin-destination pairs about three miles apart from each other. Since these are mostly main artery roads, they provide the main grid for efficient travelling between those origin-destination pairs. In contrast, minor roads and the areas between the *Oranges and Lemons* (i.e. main roads that are printed in yellow and orange in most maps) are learnt by studying the *quarter-miles* and linking the additional *points* in those areas (**Figure 3a** and **2b**). Further understanding and flexible linking is gained from the Blue Book *runs* as students start considering continuations between them. For instance, one Blue Book *run* would have continued along a sequence of straight streets, but the *run* required a turn off from this straight sequence of streets to reach a destination. In contrast to the previous example, parts of a different *run* might continue straight, where the initial *run* required to turn off the straight sequence of roads. Both examples highlight the importance of the ability to flexibly use individual *runs* as part of the ‘bigger picture’ (interview with K.T., **Appendix A**).

Ultimately, they cover large distances across London as such a combination of knowledge enables trainee drivers to link the Blue Book runs efficiently where they intersect, or through minor roads of the quarter miles where no intersection is available **(Figure 3c)**. Over time, links become more efficient as the Knowledge is ‘ingrained’ and minor roads are integrated to create shortcuts where possible. At this point, the Blue Book is no longer perceived as a list of individual routes, but as an entire network of runs (interview with K.T., **Appendix A**).

### Learning Methods

The progress that *Knowledge* students have to make from learning the first *points* and *runs* to flexibly plan routes all across London is supported through a range of learning techniques as listed in **Table 1.**. These methods can be categorized into theoretical, map-related studies and practical, ‘*in-situ*’ experiences (interview with K.T., **Appendix A**; Lordan, 2018). Both support the development of planning strategies that are later used in situations where route planning is required. These include practicing the planning of Blue Book *runs* and general *runs* with a ‘*call over partner*’ (i.e. a *Knowledge school* study partner) in preparation for exams and when driving a taxi as a qualified driver.

**Table 1.**
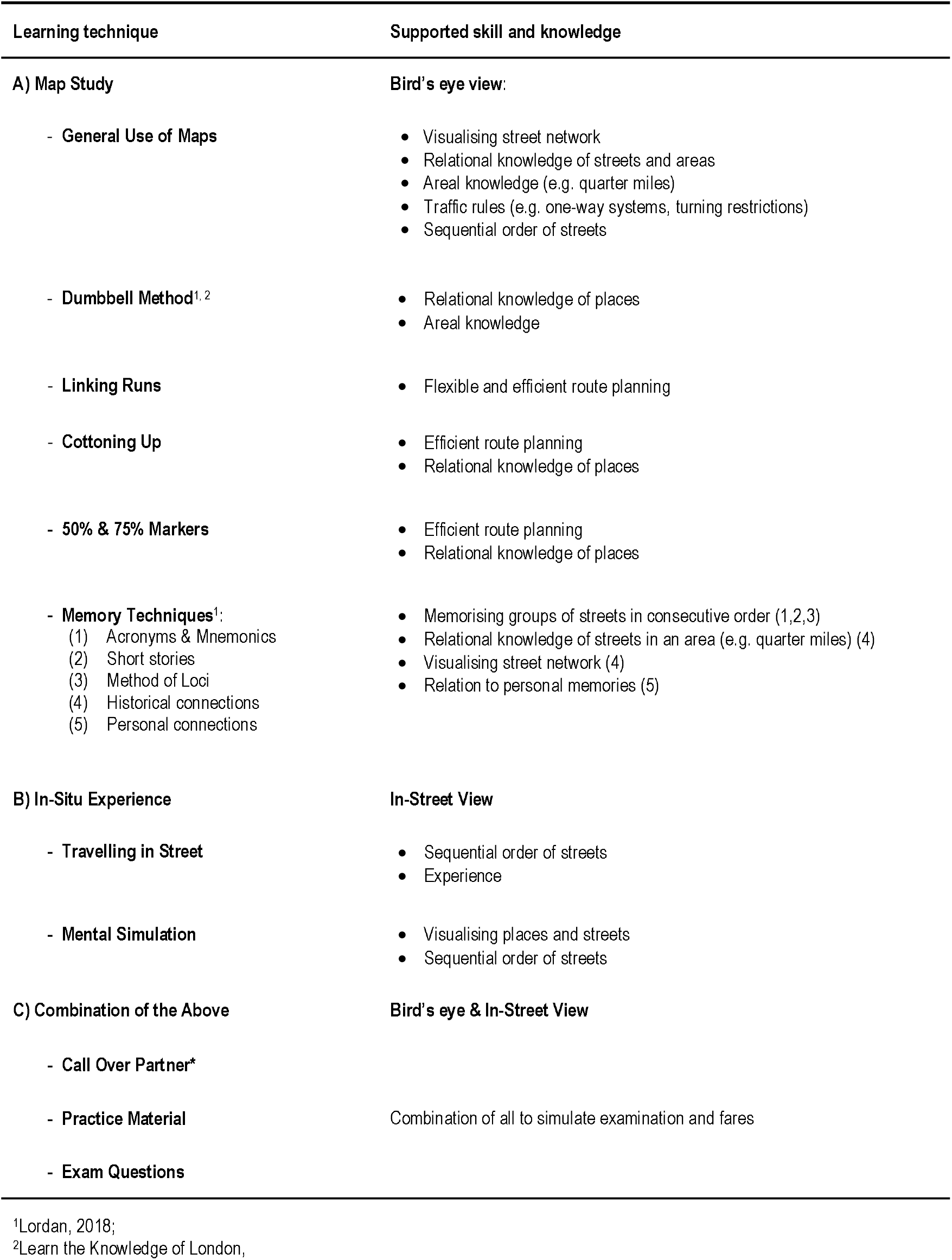
Learning techniques used in Knowledge schools.

| Learning technique | Supported skill and knowledge |
| --- | --- |
| **A) Map Study** | **Bird's eye view:** |
| - General Use of Maps | • Visualising street network<br>• Relational knowledge of streets and areas<br>• Areal knowledge (e.g. quarter miles)<br>• Traffic rules (e.g. one-way systems, turning restrictions)<br>• Sequential order of streets |
| - Dumbbell Method<sup>1, 2</sup> | • Relational knowledge of places<br>• Areal knowledge |
| - Linking Runs | • Flexible and efficient route planning |
| - Cottoning Up | • Efficient route planning<br>• Relational knowledge of places |
| - 50% & 75% Markers | • Efficient route planning<br>• Relational knowledge of places |
| - Memory Techniques<sup>1</sup>:<br>(1) Acronyms & Mnemonics<br>(2) Short stories<br>(3) Method of Loci<br>(4) Historical connections<br>(5) Personal connections | • Memorising groups of streets in consecutive order (1,2,3)<br>• Relational knowledge of streets in an area (e.g. quarter miles) (4)<br>• Visualising street network (4)<br>• Relation to personal memories (5) |
| **B) In-Situ Experience** | **In-Street View** |
| - Travelling in Street | • Sequential order of streets<br>• Experience |
| - Mental Simulation | • Visualising places and streets<br>• Sequential order of streets |
| **C) Combination of the Above** | **Bird's eye & In-Street View** |
| - Call Over Partner* |  |
| - Practice Material | Combination of all to simulate examination and fares |
| - Exam Questions |  |

<sup>1</sup>Lordan, 2018;
<sup>2</sup>Learn the Knowledge of London,

In general, maps are used to learn the structure of the street network from a *bird’s eye view*. They help obtain knowledge about relations between places and areas (e.g. quarter-miles and boroughs) and learn traffic rules that can limit route planning due to one-way systems and turning restrictions. Additionally, maps facilitate a better understanding of the sequential order of streets that are part of a *run*.

Initially, when studying the *Knowledge*, this information is obtained mainly through the ‘*dumbbell method*’. This requires students to identify the *quarter-miles* of the origin and the destination and visualise the connecting Blue Book *run* by tracing it on the map. By including variations of origins and destinations from the quarter-miles on the map, students start to connect nearby *points* with the original Blue Book origins and destinations and create a network that is forming the ‘dumbbell’ (**Figure 4**). This method is later extended to other places, as students learn to flexibly link *runs* and cover larger distances across London. This is also supported by the ‘*cottoning-up*’ and the use of subgoals, called the ‘*50% markers*’ (interview with K.T., **Appendix A**). These 50% markers (not always chosen halfway along the direct line) are bridges if the river needs to be crossed to ensure efficient planning through these bottlenecks at early stages, or other major roads and places. Additional subgoals are added before and after, as needed, to help give initial direction for the route planning without overwhelming the students. Both methods, the ‘*cottoning-up*’ and the ‘*50% markers*’, when used during initial stages of the training, help students to correctly visualise the map and relations between places. At a later stage of the *Knowledge*, when route planning is carried out mentally and without a physical map, these methods are integrated in the planning process automatically.

To help students memorize sequences of street names that are often used for *runs*, different memory techniques are applied during the learning process and often remembered years after obtaining the license. The most common techniques are creations of acronyms and mnemonics, inventions of short stories that contain street name references, mental walks through rooms of an imaginary house, historical connections and personal memories that logically structure (cf. **Table 2,** Lordan, 2018).

**Table 2.**
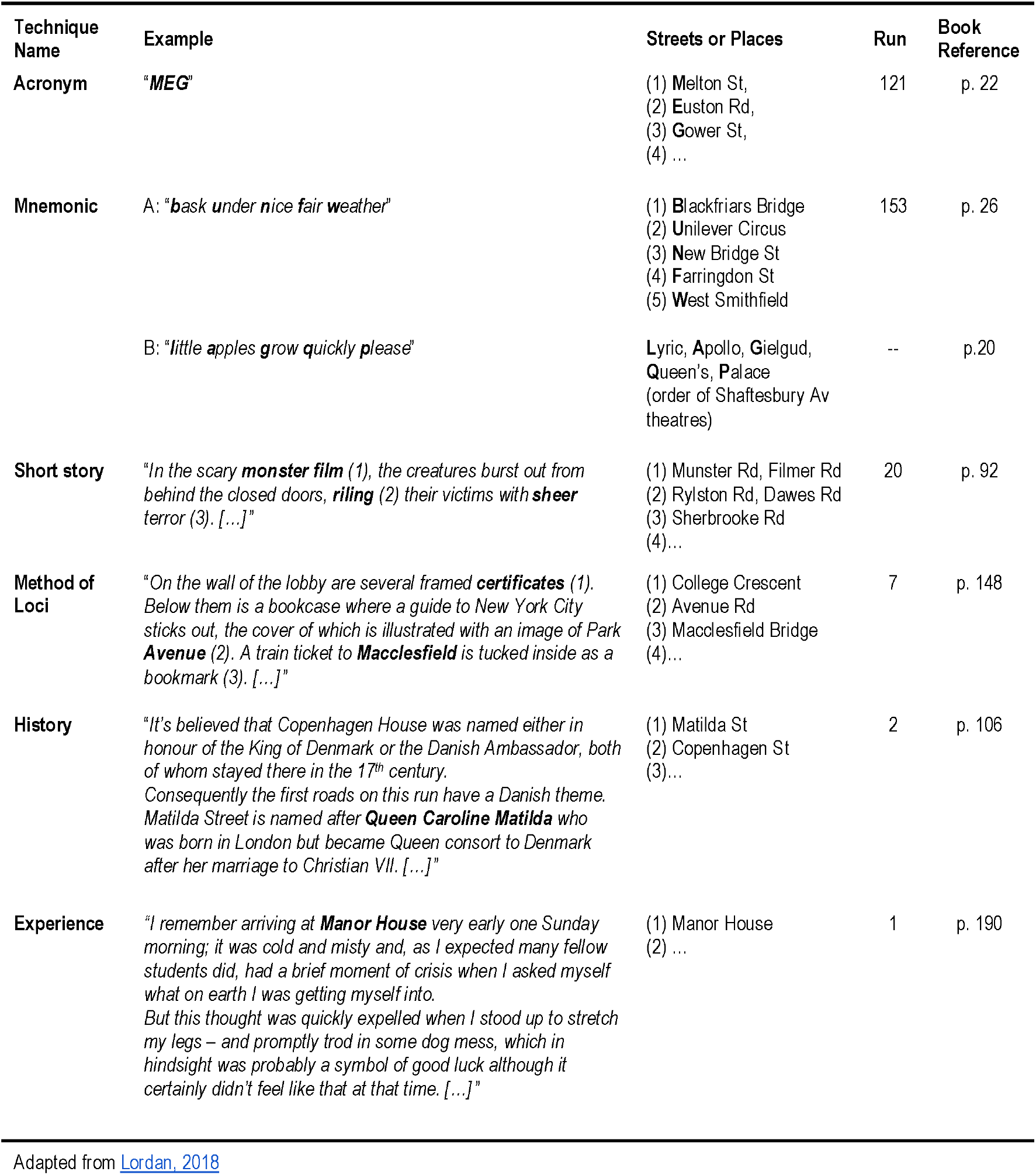
Common memory techniques to learn runs.

Location specific information from an *in-street view* is learnt through ‘*in-situ*’ visits to the 320 origin-destination pairs of the Blue Book, their *quarter-miles* and driving the corresponding *runs*. These visits - carried out multiple times, often on a scooter with a map of the Blue Book *run* attached to the windscreen - are essential to learning and recalling the *Knowledge*. These experiences of runs and the quarter miles create memories that drivers use to later recall sequences of streets (**Table 2,** Lordan, 2018) and visualise routes during planning (interview with K.T., **Appendix A**). For instance, memories of travelling a run for the first time might help the recall of sequences of streets, places of interest and specific traffic rules that must be obeyed. These memories become an essential source of information when planning and calling out similar *runs*, linked to the original. Students use them for mental simulations that facilitate decisions about where to pick up or set down passengers, in which direction to leave or to approach an area and how to find the most optimal route.

### Assessment Scheme

The assessment scheme for trainee taxi drivers in London was designed to support the learning process and guide students from early stages of learning the initial Blue Book *runs* to final stages, where their knowledge of London and suburban artery roads is rigorously challenged (**Figure 6**; interview with K.T., **Appendix A**, Learn the Knowledge of London, Transport for London, n.d). Initially, *Knowledge schools* offer an introductory class to provide basic information and an overview of the content of the *Knowledge*. This introductory class includes expectations, procedures and requirements of the qualification process, before preparatory examinations (**Figure 6**, light grey) can be taken. Within the first six months of starting the *Knowledge*, students are expected to sit an assessment that is testing the *Knowledge* on the initial 80 *runs* (five lists) of the Blue Book. Even though this assessment is unmarked, it is obligatory and of supportive and informative purpose at the same time (i.e. formative assessment). Feedback is given and the performance is discussed with teachers to help students identify problems in their learning process that need adjustment at an early stage to enable students to successfully progress at later stages. Following this initial self-assessment, students have 18 months to sit a marked multiple-choice exam that tests their knowledge of the Blue Book, to ensure they have acquired the basics that are necessary to progress to the appearance stages (**Figure 6**, dark grey). To test this, the multiple-choice exams consist of two parts, where (a) the shortest, legal route out of three possibilities has to be identified for 5 randomly chosen Blue Book *runs*, and (b) the correct location out of six possible locations has to be selected for 25 points of interest that are likely to be part of the learning of the Blue Book *runs*.

**Figure 6.**
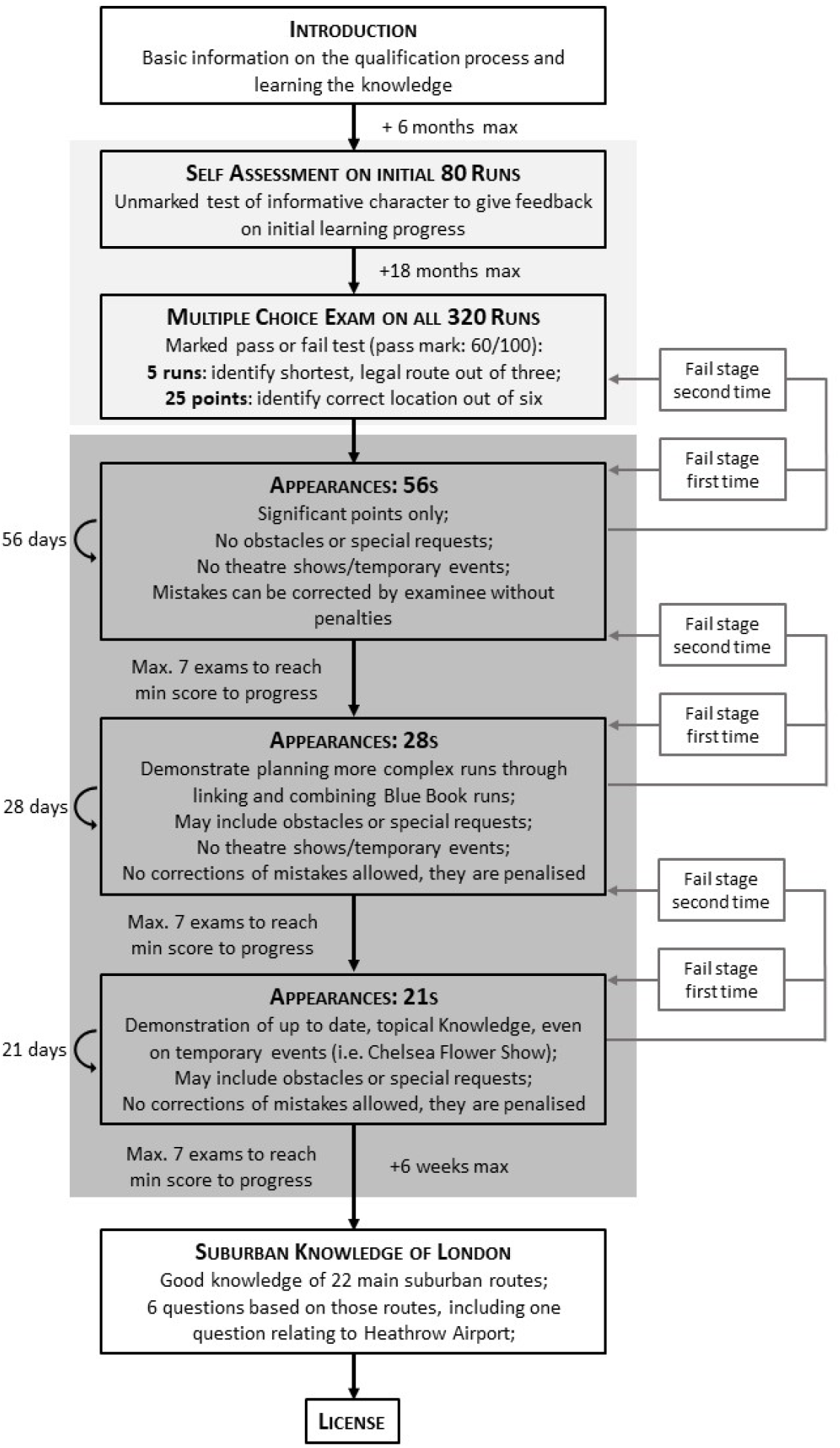
Knowledge examination process. The initial stage (light grey) of the *Knowledge* examination process provides feedback (*Self Assessment*) on the individual progress of learning the first 80 *runs* of the Blue Book and assesses the minimum knowledge on all 320 Blue Book *runs* needed (*Multiple Choice Exam*) to start the oral examination (*Appearances*). The main part of the examination process (dark grey) consists of a series of oral examinations, the so-called ‘appearances’, consisting of three different stages (the 56s, 28s and 21s, named after the intervals between each exam in the corresponding stage). Even though the requirements to students sitting these exams become more rigorous as they proceed, there are general rules that apply across all stages. These are related to the general layout of each appearance (e.g. duration, number of *runs*), expectations (e.g. shortest route), format of call out (e.g. identifying the location of origin and destination, sequentially naming streets and providing turning instructions), penalties (e.g. traffic rule violations, deviations from shortest route, hesitations), awarded points and progressing to the next stage. Following the appearances, students are required to pass an exam on suburban *Knowledge* before they obtain their license. Source: Adapted from <u>Learn the Knowledge of London</u>; <u>Knowledge of London learning and examination process</u>, p. 21

After passing the two entry assessments, trainee taxi drivers enter what is known as the ‘*appearances*’, a set of oral examinations. At each *appearance*, students are expected to call *runs* from any two *points* that the examiner names. The *appearances* also comprise the longest and most difficult part of the *Knowledge* examination process. It is quite common that several of the stages have to be re-taken by students due to shorter intervals between appearances coupled with the growing expectations of the examiners as they proceed. In total, there are three stages of appearances, the 56s, 28s and 21s, which correspond to the number of days between any two appearances in that stage.

Even though the requirements for students sitting these exams become more rigorous as they proceed, there are general rules that apply across all stages: Each appearance is about 20 min long and can consist of up to 4 *runs* that students have to call, using the shortest route, disregarding traffic and temporary roadworks. The *call outs* (i.e. naming streets in sequential order) include identifying the location (i.e. the correct street) of the origin and destination (points of interest), naming streets and giving turning directions along the *run* in correct sequential order, as well as including instructions for leaving and setting down passengers. Possible errors that will cause deductions of points are incorrect street names, any divergence from the shortest route, violation of traffic rules, impossible leaving or setting down instructions and hesitations during the call of the *run*. In each appearance, 3-6 points are awarded and 12 points are needed to progress to the next stage. Per stage, students are allowed to fail a maximum of three appearances, before the stage has to be repeated (first time) or students have to go back to a previously successfully passed stage (failing second time), limiting the number of exams per stage to a maximum of 7 appearances.

In contrast to later appearance stages, the ‘56s’ are very closely related to the *Knowledge* obtained from the Blue Book. Here, examiners closely stick to *runs* from the Blue Book, which reflects a good knowledge of primary and secondary roads (i.e. the ‘*oranges and lemons*’). At this stage, examiners also take into account differences in the choice of additional *points* of the quarter-miles that different *Knowledge schools* provide in their version of the Blue Book (**Figure 3a**). Additionally, *runs* are structured in a way that they will not contain obstacles (e.g. road closures), special requirements (e.g. requests to avoid traffic lights) or theatre shows and temporary events (e.g. Chelsea Flower Show). Students are also allowed to correct mistakes by going back in their call out and changing their *run*. At the next stage, the ‘28s’, examinees are expected to be able to link *runs*, using some minor roads and avoid obstacles or comply with special requests without being granted a chance of correcting faulty *runs*. At the final stage, the 21s, trainee drivers have to demonstrate an overarching knowledge that is up to date and can additionally refer to particular topics (e.g. new tourist attractions, changes in hotel names) and temporary events, such as the Chelsea Flower Show.

After passing all appearances, the final exam is set to test the knowledge of suburban London. This knowledge covers 22 specific routes, including major *points* along those routes, radiating from the six-mile radius to the borough boundaries of London. In this final appearance, trainee drivers will be asked six questions relating to the 22 routes and *points* along those routes.

For the learning process of a *Knowledge* student, the Blue Book is central, as it provides them with “*the ability to know where streets and roads are going to and where all those places are*” (interview with K.T., **Appendix A**). However, over the course of obtaining the *Knowledge* and learning how to link Blue Book *runs* efficiently, there seems to be a change in the perception of London. Initially it consists of distinct routes and locally focused areas on a map. Over the course of time, this fades into a connected, large-scale, inseparable network of streets and places in the real world (Appendix A). During consulting conversations with taxi drivers, they reported that they just knew where they had to go without much planning. For well-known places, Robert Lordan described the planning and execution of a *run* as “*I wouldn’t even have to think; my brain would be on autopilot. […] like a moth drawn to a light!*” (email conversation with Robert Lordan, **Appendix B**). For longer distances, subgoals (as trained with the 50% markers) are used automatically: “*I’d find that my brain would often plan in stages; essentially I’d envision a set of waypoints and the route would then come to me as I progressed*” (email conversation with Robert Lordan, **Appendix B**).

The overall impact of the *Knowledge* also seems to foster a deeper connection (“*I already loved the city, but in studying it I now love it all the more. It feels like an old, familiar friend*”, email conversation with Robert Lordan, **Appendix B**). It provides a constant drive to stay up to date with changes in the city (“*The Knowledge made me crave detail! To this day I want to know as much as I can about London*”, email conversation with Robert Lordan, **Appendix B**) and new curiosity (“*The Knowledge also makes you want to know as much as you can about new locations that you’ve never been to before*”, email conversation with Robert Lordan, **Appendix B**).

## Discussion

Here we examined the process by which licensed London taxi drivers learn and are examined on the Knowledge of London, which includes the network of ∼26,000 streets and thousands of points of interest. In summary, to learn the Knowledge of London, taxi drivers use a wide range of theoretical and practical methods and learn specific methods for efficient planning. Such training primarily includes map-related study, based on an overlapping network of basic points of interest and list of routes (Blue Book) that systematically covers London. This knowledge is combined with visits to the locations used in the routes and retracing of the theoretically learnt routes on motorbikes. Both experiences are reported to be vital for linking theoretically learned information to specific real-world locations and flexible navigation in London. We also observed a range of techniques to improve memory, such as acronyms and stories linked to sequences of streets, visualising the locations and travel along streets, and the strategic use of sub-goals. We discuss: i) how these findings relate to other studies examining spatial learning, ii) how the learning compares with taxi drivers in other cities, iii) why the knowledge is still required and trained when GPS aided navigation systems exist and, iv) how these methods and techniques might benefit the general population in spatial learning.

Research based studies of spatial navigation have employed a variety of methods to train participants learning unfamiliar environments. These include instructed learning of paths (e.g. Brunec et al., 2020; Wiener et al., 2013; Meilinger, Frankenstein et al., 2014; Meilinger, Rieke et al., 2014; Meilinger et al., 2008), learning from cartographic maps (e.g. Coutrot, Silva et al., 2018; Coutrot et al., 2019; Grison et al., 2017; Holscher et al., 2006; Holscher et al., 2009), landmark-based navigation (e.g. Wiener & Mallot, 2003; Wiener et al., 2004; Wiener et al., 2012; Wiener et al., 2013; Astur et al., 2005; Newman et al., 2007), exploration of the environment without a map (e.g. Spiers et al., 2001; Spiers et al., 2001b; Hartley et al., 2003; de Cothi et al., 2020) or a combination of map study with in-situ exploration (e.g. Javadi et al., 2017; Javadi et al., 2019a; Javadi et al., 2019b; Patai et al., 2019; Wiener & Mallot, 2003; Wiener et al., 2004; Spriggs et al., 2018; Warren et al., 2017; Newman et al., 2007). The general assumption is that the method used for learning is efficient, or a standard way of learning the environment. Here we found that for London taxi drivers the training is significantly more intensive and elaborate than any of these studies, which relates to the dramatically increased demands of learning 26,000 streets and thousands of points of interest. Several methods for learning, such as guided turn-based navigation (e.g. Wiener et al., 2013), have not found an application in the training phase of London taxi drivers. The absence of this approach might be explained through the advantage of in situ experience, understanding the changes with lighting over day time and the very regular changes to the environment (e.g. temporary road closures, name changes of hotels or restaurants, and temporary events). Indeed, being able to adapt to these changes and being aware of some of the temporary events are considered essential knowledge, especially at later stages of the training process.

Successfully recalling mental images of locations, retrieving specific street names and judicious uses of sub-goal planning were described as key to being a London taxi driver. These observations help to explain results of by Spiers and Maguire (2008) where London taxi drivers were asked to recall their thoughts watching video replay of their navigation of a highly detailed virtual reality simulation of London. London taxi drivers often reported sequential planning to sub-goals along the route, comparison of route alternatives or mental visualisations of places and route sequences. Many taxi drivers reported ‘picturing the destination’, planning with a bird’s eye view, and ‘filling-in’ the plan as they navigated, which indicate a use of mental visualisation as trained through the Knowledge. We found teachers and examiners claim to know when students ‘see the points’ as they actively visualise origins and destinations as part of their planning process. It may be that trainee taxi drivers need some ability with mental imagery to succeed in the train process. Not all trainees will pass the examination process (Woollett and Maguire, 2011). The ability to use spatial visualization strategies has been found to differ between individuals and vary with age and experience (Salthouse et al., 1990), education levels or gender differences (e.g. Wolbers & Hegarty, 2010; Coluccia & Louse, 2004; Montello et al., 1999; Moffat et al., 1998; Fennema & Sherman, 1977). There is also evidence that certain spatial visualisation skills can be improved through training (Sorby, 2009). In our study we found that it was expected that the visualisation improves with the training. Further investigation of the visualisation process in novice trainees and expert drivers would be useful and may relate to the changes in the hippocampus observed in those that past the exam to obtain a licence (Woollett and Maguire, 2011).

Further evidenced use of mental simulation during navigation was found in the way taxi drivers are required to call out the runs in the exam by using instructions and phrases such as ‘forward’, ‘left/right into’ and ‘comply’ (traffic rules). These provide an egocentric description of movement through London. Conversely, during the early stages of the Knowledge training, the planning process is reported to rely on an allocentric reference frame by studying maps to train students on planning shortest paths. At later stages, as experience is gained from planning runs and through in-situ visits to locations, the aim is to build an automatic awareness of the direction of travel or a particular route. This is consistent with the reports that experienced taxi drivers very rapidly determined the direction to a requested destination (Spiers and Maguire, 2006; Spiers & Maguire, 2008).

We found that the examination process appears to provide a layered approach to learning the London street network. There is an initial focus on testing the Blue Book routes (runs) or routes along main arterial roads (i.e. ‘oranges and lemons’) and only at later stages are minor roads integrated into the assessments. However, we found the actual learning process requires students to learn minor roads in the quarter-miles from the beginning (i.e. with the first run). This differs from the requirements in other cities, such as Paris, where drivers have to demonstrate knowledge of a limited number of major points of interest, as well as predefined major routes. There, taxi drivers are expected to expand their knowledge to the minor street network through experience whilst working as a taxi driver (Prefecture de Police, Demarches & Services, 2020; Skok, 2004). Similar to the ‘oranges and lemons’ of the London street network, the Parisian street network covers the city in two layers: The base network, an uneven grid-like pattern that allows travel on major roads, helps to reduce traffic on the secondary network, a network of minor streets (Pailhous, 1969, 1984; Chase, 1982). For Parisian taxi drivers, such a selective learning of the base network was found to be also reflected in their mental representation of the street network in form of these two layers (Pailhous, 1969, 1984). In contrast to London taxi drivers, Parisian taxi drivers’ awareness of the secondary network only grows and becomes more efficient and optimal through experience rather than in the training and is almost non-existent at the beginning of their career (Chase, 1982; Giraudo & Peruch, 1988, Peruch et al., 1989).

The approach that London has taken to train and test their taxi drivers on the Knowledge as described above, is historically motivated and has been retained over centuries since its implementation, only allowing for adaptations and improvements. This concept of learning all possible points, their locations, the street names and how to flexibly plan routes and adjust to specific requirements is globally unique. In contrast, other cities, such as Paris (Prefecture de Police, Demarches & Services, 2002) or Madrid (Federación Profesional del Taxi de Madrid: Departamento de Formación, 2010; Skok & Martinez, 2010), often only require applicants of the trait to learn the major grid of the street network (i.e. the base network) and expect the knowledge of the minor street network (i.e. the secondary network) to be obtained through experience. Instead, taxi drivers are also required to demonstrate knowledge on other trade related areas, such as knowledge related to driving a car, professional regulations, safety and business management, a language test (Skok, 2004), fares, legislations and personality (Skok & Martinez, 2010). Considering these alternative qualification requirements for Paris or Madrid, the London qualification scheme, that relies on a thorough knowledge of London streets, can be questioned as regards to its adequacy and value, in times of GPS systems that can guide navigation.

Given that GPS in general successfully supports navigation and thus is omnipresent in daily life, it remains a key question as to why London taxi drivers continue to rely on their own abilities to plan routes. We found that this to be their sense of accomplishment of a difficult, and in this case, almost impossible task. They often find pride in their ability to master challenging navigation tasks in a complex city only by using their spatial memory independently from external devices that could be sources of mistakes (McKinlay, 2016). This ability to flexibly navigate beyond a base network of major streets, enables London taxi drivers to rapidly follow their route plan even to points in the secondary network, quickly adapt to any changes on-route due to customer preferences or traffic flow (i.e. congestion or road closures) and avoid errors that might result from incorrect instructions given by passengers (e.g. Lordan, 2018). For instance, they might confuse Chelsea’s buzzing shopping mile, King’s Road, with the quiet King Street near St James’s Park, Westminster. These adaptations, that taxi drivers can make instantly, might even outperform GPS systems that sometimes need manual adjustments and additional information input to get to a similar result. In contrast to London, it takes taxi drivers in Paris, Madrid and other cities years to acquire this type of knowledge in their cities and in the end they might never achieve a similar, highly accurate knowledge of their cities as some areas might be less frequently travelled. Moreover, their experience to filling the gaps in their knowledge might strongly rely on their use of GPS devices, which have been found to impair spatial learning (e.g. Ishikawa, Fujiwara, et al., 2008) and interfere with spatial navigation (McKinlay, 2016; Johnson et al., 2008). These methods of training taxi drivers might be less efficient and it is thus not surprising that there have been requests from taxi trades of cities like Tokyo, asking London Knowledge teachers to develop a similar method for their taxi schools (interview with K.T., Appendix A).

How might the Knowledge training process be improved? The Knowledge in its current form, based on the 320 Blue Book runs, has been in place for about two decades, but the study methods have remained the same over many more decades. However, there has been a tendency of involving new technologies and creating online resources, such as apps that can hold and test students on the Blue Book runs. By providing the first plot of all the blue book runs we were able to identify regions in the road network that were poorly sampled and it may be possible for this information to be useful should new routes be required in updating the runs.

It is possible that a data base of videos of Blue Book runs would be useful. However, updating this database is a challenge due to the regular change in London’s appearance and layout. Online maps and applications could provide a platform that could be regularly updated. Here, the focus could be on Knowledge requirements that allow general contribution, similar to OpenStreetMaps, and individual modification, as with Google My Maps (Google Maps. My Maps, n.d.), to support the individual learning process. Such a platform could include updates on points asked in recent appearances that students use for preparation or an option to train with and challenge other students, as well as their call-over partner. However, these platforms would not be able to replace the social situations that students find themselves in at Knowledge schools and when practicing face to face with their call-over partners. These social interactions also have a psychologically motivating, supportive effect. Neither can these digital maps overcome some obvious visual limitations due to screen sizes. These will not allow for a similar view of the ‘bigger picture’ that a wall-paper map is able to convey.

How might the learning process described here be exploited for the general population to learn new places? A number of recommendations could be made. One is the focus on street-names. Much navigation in cities can be based on landmarks and the rough knowledge of the area, but by learning the specific street names it increases the capacity to consider routes between locations and understand the inter-relations in the space. This learning can be enhanced by a focus on methods to draw out the street names such as acronyms and rhymes (‘East to West Embankment Best’). The memory techniques used in Knowledge schools to memorise sequences of streets such as the ‘dumbbell method’ that links small areas through routes, or mental visualisations of familiar places could initiate new ways of displaying spatial information in maps or GPS devices. A focus on mental imagery is also worth considering in future research to explore how this may benefit new navigation. Finally, teaching a method for efficient planning of longer routes would be a benefit. More research will be required to fully explore these possibilities and understand how they may be integrated with other technology for efficient spatial learning.

Another question arising is how might these discoveries be useful for researchers seeking to build efficient AI systems capable of rapid learning and planning? The main discoveries here that may be relevant are: 1) the organised learning of a set of interconnected routes that allows for flexible planning in the future, 2) the focus on learning a route and then exploring the points at the start and end and then connecting the route to other routes, 3) learning to create sub-goals during the planning process. These approaches to learning may extend not just to improving guidance for how humans learn but for considering the construction of agents that optimally learn structures in the layout of a large city network.

In conclusion, studying the training process of licensed London taxi drivers has provided a useful opportunity to better understand learning strategies and methods that efficiently support the learning process of a large and complex environment. In this observational report, information was gathered on licensed London taxi drivers, who acquire unique spatial knowledge to navigate an enormous street network independently from external support, such as GPS. Forming such mental representations of real-world spaces is essential and training strategies in experimental settings vary. Essential strategies include memory techniques, map-based strategies using tactical subgoal selection to improve planning efficiency and mental visualisation of places and routes based on experiences. Further research is needed to understand the mental representation that results from these training methods and how this representation affects navigation related planning in brain circuits including the hippocampus.

## APPENDIX A

Interview with a teacher from a Knowledge School in London, UK, about the Blue Book Int: Interviewer

KT: Knowledge School Teacher

Note: Hesitation markers (e.g. uhm) have been removed from the transcription.

Int: I’m interested in things about… anything that you know about the Blue Book. And how it developed, a bit of history, why it is structured the way it is structured. If you could just tell me then I’ll probably ask whenever there’s a lack. (0:21)

KT: Well, the Blue Book, as it’s called, has been around for over a hundred years. But it has developed. In the year 2000, it was completely redesigned. London had changed dramatically. Some parts of it had not really been covered on the old Knowledge. So, a gentleman from the Carriage Office was tasked to completely redesign the Knowledge. So, what he did is he set about dividing London up into learnable pieces, like the small jigsaw pieces. And after consultation, he realised that a quarter of a mile area was the ideal amount that a person could easily absorb. (1:09)

Int: Was he like a taxi driver himself?

KT: No, he had been, but was an ex-police officer and he was also an examiner at the Carriage Office. What he did was, he divided London up into quarter miles. And when he’d finished this circle, there ended up being three hundred…, six hundred and forty of them. (1:32)

Int: Ok, so he developed the circles across London first and then he decided on connecting them up…

KT: Connecting comes afterwards. Now, from my own personal perspective, I have been to Tokyo with the Japanese taxi corporation. They asked me to show them how they could develop a Knowledge. I also… I turned the opportunity down… I could have gone to Korea and done exactly the same thing. So, what he developed would work in any major city. And that’s the thing. I don’t think a lot of people realise. (2:00)

[*parts omitted to keep anonymity*]

KT: It was in the year 2000. Effectively what he did, when he’d finished dividing it up in these bite size pieces, now he did consult with us, and we did say that a quarter mile was just about right to learn. And when he’d finished covering London in those quarter mile circles, coincidentally, it amounted to 640. Then what he had to set about doing, was making certain that each circle had an overlay with a neighbouring circle. Then he had to ensure, and I will show you some examples in a minute… he then had to ensure that you would leave an area in each direction. So, if we start at Manor House, the first run leaves Manor House and heads towards Gibson Square. So, you’d learn that area of a quarter of a mile around Manor House and you exit the area heading south. Now later on, you come back to the edge of that area to a place call Harringay Green Lanes Station, but you arrive there coming from the north. Is that making sense? (3:17)

Int: Yes, it does.

KT: Later on, there is several other runs in the area and they all leave the area in different directions. Now, the more able candidate will realize this very quickly and he will be able to link together. The less able candidate sees the journeys as individual journeys and fails to make a connection at the start and the end of each run. Is that making sense? (3:46)

Int: Yes.

KT: Now, within each quarter mile, the candidate is expected to learn key places of interest. Now, obviously in the centre of London, there could be as many as forty to fifty places within a quarter mile that need to be learnt. As we move further out to the edges of the six-mile radius, there will be less key points of interest. Now, key point of interest it’s pretty obvious. It will always be a hotel, a restaurant, a theatre, large restaurants, religious establishments, anything that the public will need to go to, the candidate is expected to know. (4:25)

Int: You said closer to the centre there are more places than… like more places within the quarter mile radius, rather than further out. But in the Blue Book there are usually about ten.

KT: Right, well that… I’m afraid, I was responsible for that. There is actually eight. And that was again as a result of teaching students and looking at what information they could hold. And we realised that if you gave them more than ten, or more than eight even, they gave up. So, what we decided to do, was to make it a standard eight, as was part of the learning process. Later on, of course, they will then access material which will give them more points in key areas. So, west one (W1) which is the central area of Westminster, will have a lot more points in it, than say E ten (E10), which later, which is at the edge of the map. (5:20)

Int: Ok, but these additional points they learn later? KT: They will find those at a later stage.

Int: And those eight key points, they are the same for every single knowledge school, (…)?

KT: No, no, no… Knowledge Schools develop their own training material. I developed them based on what examiners asked in the past and obviously on my knowledge of what I felt were the most crucial points within the quarter mile radius of the start and the end of each run. Now, if we move on to the runs, the idea of them is that on a map of London, the roads are coloured orange and yellow. Orange for the major roads, yellow for the secondary roads. The idea of them is that each quarter mile was linked to another one, approximately two to three miles away. And that gave the candidate an opportunity to drive along the orange roads or the yellow roads. The smaller roads, should have been found and learnt at the start and the end of each run. (6:20)

Int: Okay,

KT: Is that making sense? Int: Yeah, that makes sense.

KT: Now, if you come over here, that effectively is what I’ve just told you.

*[shows map of London with quarter miles, see Figure 2]*

KT: If you look at that, that is all of the beginnings and the endings of the runs. If you look at the map closely, you will notice the orange roads and the yellow roads. And effectively all of those roads are covered in the runs, as well as the significant grey roads, which are lesser roads. (6:51)

Int: Okay, do you cabbies have to learn all of the white ones as well? KT: In theory, yes, in practice, no, it would be impossible.

Int: Okay, I heard from previous lessons, that I mean, if you look at them, the first 80 make up like a first layer…

KT: That’s exactly it, yeah…

Int: …and then a second layer, and then a third layer, and… KT: I can actually show you the first 80.

Int: Why was it chosen as 80 – 80 – 80 – 80? Like four times 80?

KT: That wasn’t. That was us, that was aesthetics. That was the knowledge school dividing it into that system. That’s what I’ve been looking to show you. So, the first 80 for example, that’s how they cover. (7:37)

Int: Ah, okay.

KT: Is that making sense?

Int: Yeah.

KT: When you come to the second 80, you’ll find that they will start next, and then the third 80 and the fourth 80 and by the time they’re finished, there are no grey areas left. So, learning the smaller roads, should really be done, when you’re doing your orbit and your quarter mile at the start and the end. Now the course that I developed, the Compass Direction Course, one of the things that I found candidates have a problem with, they learn the roads, they learn where they’re going to, but when you take the map away from them, they then have to see the map without it being there. Does that make sense? (8:14)

Int: Yeah.

KT: So, in other words, if you got into my taxi here, outside this building we’re in here now, and you said you want to go to Wandsworth, immediately, I have to plot the route, it’s not about me knowing the road, but I have to understand the route. So, I developed a system, what’s called bullets. So, when the run goes across the River Thames, the 50% bullet would always be the bridge. So, having as attained for our example I would choose Albert Bridge from here. I then got to get myself very quickly something equidistant between here and Albert Bridge, a target, ich which case it would be Trafalgar Square. So, my mind says, right, drive to Trafalgar Square, drive to Albert Bridge, and then I’ll make the journey much easier for myself, because as I come across the bridge, I can then focus it on Wandsworth. Is that making sense? (8:58)

Int: Yes, makes sense. How are these bullet points connected to the, or are they at all connected to the Blue Book runs?

KT: No, not really, because they’re done at an advanced stage. The candidate that will come to do a Compass Direction Course, which I developed, will only come to me having completed all the Blue Book runs, will probably have a substantial knowledge of the points in and around the area. What they will tend to need is this guidance of how to quickly make your mind up about a journey across the top of the map, diagonally down the map or which ever. (9:34)

Int: Do you also use these bullet points for shorter routes?

KT: Not really, no. The shorter runs come just linking up… I’m doing a class tonight with them. The shorter runs, come through linking up the quarter miles, so driving through a few quarter miles.

Int: Okay. So, it’s mainly for the long runs.

KT: It’s mainly for the longer runs where you have to use bullet points. Int: And you said you have 50%, usually the river, and then in between…

KT: What I get them to do is, if you sit down and try and think of a journey in one go, it just… it’s too much. So, what I say to them is, quick, quickly think of which direction you are going to heading, and then say to yourself, all right, on the map, I’ve got to go towards that place, be a junction, a roundabout, a bridge, whatever. So, get me there. Then you’ve reduced the journey in your mind that you’ve got to view. Does that make sense? (10:32)

Int: Yeah.

KT: So, when you’re at the half way mark, then it’s much easier to think, ah, yeah, I’ve got to head towards there now.

Int: Okay. So, coming back to the Blue Book, and the map. So, you said like you started off with Manor House, and then you go down to Gibson Square, from there you leave…

KT:…You link the next run, which is Thornhill Square. Int: Okay, how far is that away?

KT: Usually they are less than a quarter mile away. (11:04)

Int: Okay, and those quarter mile radiuses then overlap in the beginning and that’s where the linking happens.

KT: Absolutely, and that’s where you start to learn the backroads by going from the end of one run to the start of the next. That helps you connect it. (11:19)

Int: So, if you would draw all the runs, one after the other, is there like some kind of system like for the first 80, because the first 80 roughly overlap.

KT: No. That’s a good question. They are quite random. I have done it to see. But what they do do, they introduce you to all the key orange roads. Although you don’t travel along the whole length of that road. What I used to encourage those beginning students to do, was to draw the run on the map, but where you leave a major orange road, get a red pen and just see where you left it to where it would have continued on to. So, one long run for example, number two I think, they came down into King’s Cross Road, and then the run took them up a street, called Acton Street, so I used to stop the class to say, right, now with the red pen, I just want you to look to see, although you turn off that major road, I’d like you to understand where it would have gone to. Because later on, they have another run that meets that road further down. Does that make sense? (12:28)

Int: Yes,…

KT: So, I’m trying to get them to connect this bit of it. They leave it here and then they come back at onto it here. But some of them, the lesser able students, don’t realise that this road, that they are coming back onto here, was the same road they travelled along up there, because it might have changed its name.

Int: Okay, okay. You said the Blue Book actually, it was like 2000, when they…

KT: It was in the year 2000, when it was completely redesigned and brought up to date. Int: Okay, and I think they reduced the number of runs… (13:02)

KT: They reduced the number of runs from 480 to 320, but that really was immature, because the lot of the runs were just duplicates. They went exactly the same way. What they… Some of them started in areas that… just give me an example… Belgravia for example, about 30 of them started or ended there, but I think Greenwich had one. So, there is a much more equal spread that you see there. (13:28)

Int: Are there any areas in your experience, any areas that kind of students find more difficult to learn or struggling more with or less, because maybe the blue book runs don’t cover those areas well or whatever the reasons?

KT: No, whoever designed this, did an incredibly good job. If you look, the only gaps that are there are parks. If you look, you can see all those circles and the pins. The only space that’s left is the green parks. What does happen, some areas because of the complexity of the road layout students do find harder. Psychologically, what has always been interesting to me, if a guy lives north of the river, he tends to worry more about the south of the river, and conversely, the guys that live south, worry about the north, if you live east, you tend to worry about the west. That’s a natural phenomenon. (14:22)

Int: Okay, but there is not one area where you can say, this is a very complicated, difficult area?

KT: Well, obviously Westend, central London and the City is always complicated. The City, because of the number of one-way streets that are there and the restrictions imposed upon them. Whereas as you get further out to the edge of the map, there’s less one-way streets and less complexity.

Int: But I think wouldn’t that be also more difficult for students to learn because it’s further out and they don’t tend to go there that often.

KT: Well, it’s harder for them to pick up points. That I will accept. I think the learning the (…) and street around the area is not too difficult, because if we look at that top run up there, 170 I think it is, there isn’t an awful lot there. It’s a main road, that takes them back in. And most of the stuff there is industrial, so I would encourage them to pick up four of five points, there is a train station at (…) road, but apart from that, there isn’t a lot there. And if we look in the centre, can you see all those circles where they’re overlapping them? That’s very, very complex. (15:26)

Int: Okay, okay. What would you improve about this if you had something to, or someone asked you? What would you do?

KT: The only thing I think I would improve would be the examination process. I think, I would probably go back to what they did try once. Where rather than let everybody have learn everything in one go, I might say to them, right, I wanna you come in after you’ve learnt 40. 40 runs, that’s 80 beginnings and endings and if you haven’t done them adequately, at that point we can give you corrective assessment. Unfortunately, the examining body doesn’t have the responsibility to correct a student and give them guidance. That’s where the knowledge school should come into the, you know, formulation, but it doesn’t always happen. In terms of the layout and learning, no, I think it’s, as I say, I’m full of admiration for the man who came up with this. (16:34)

[…]

KT: Everything I teach, every lesson I do, and I have done for, I mean I’ve been teaching the Knowledge for now over 20 years, I’ve always related everything to the Blue Book. Tonight’s class, they’ll be doing east London with me. And I’ve listed about 20 Blue Book runs. I’d expect them to be able to give me at least four to six points at the start at each of those runs. But then, what I’d expect them to do to cross connect them. So, for example, Manor House run goes to Gibson Square, but with an advanced student, I would expect them to get me from Manor House station beyond Gibson Square, say to Thornhill Square. So he can work out how the run almost is the same, but then it differentiates towards the end. (17:23)

Int: When you, when you actually do call some runs or plan runs, do you actually use lots of the Blue Book knowledge, linking up the Blue Book runs mainly rather than…? Or does it become more flexible?

KT: Yes, as they get more advanced, it does become more flexible. But certainly, the early candidates, those that were on a 56 day standard, I encourage them to stick as closely as possible to the Blue Book. Because I think the level of Knowledge they’ve got isn’t sufficient for them to always make their own decisions. (17:54)

Int: But later on, let’s say at the 21s,…

KT: Obviously, on the 21s it’s got to be random, completely random, because they have to be almost a taxi driver. And in 30 years of driving a taxi I think I only ever had two journeys that were remotely Blue Book runs. (18:10)

Int: Oh, really?

KT: Yeah, yeah. Most of the time it’s just completely random journeys.

Int: So, would you say that later on, when you really drive a cab, the Blue Book doesn’t influence you that much in your driving.

KT: No, it doesn’t but what it has done of course, it’s given you the ability to know where the streets and roads are going to and where all those places are. But a lot of the journeys that you will then undertake, some of the things you’ve learnt, will just stay on the back shelve. Simply because of time. Remember, the Blue Book is always done in the straightest line. Sometimes that’s not practical. (18:44)

Int: (…) it’s very interesting how the Blue Book is constructed, because for us it looks like a big mess. We don’t see all those structures that you have (…), but if you would think about alternative methods of learning the Blue Book, let’s say you start off in the north west and you learn every single street around that area and then you expand gradually. Or you start off in the centre of London, let’s say Trafalgar Square, and then you just expand your circle of learning the map more or less. Would that make any sense for you? It’s a different way of learning. (19:29)

KT: It is. And I’m always open to different methods, but I’m so convinced that this works. I would be reluctant to have a system that altered that. I think the danger then starts becoming that people then pick and choose which areas they want to learn. You’re with me. And it would be a great temptation to ignore these bits down south and up the north there and people would go to what we call the honey pot, which is the centre. I think they’d spend more time learning that. The idea of the blue book is that it makes you to all these far flung places. (20:02)

Int: So, it would be more flexible, with the Blue Book you’re more flexible KT: Yes.

## APPENDIX B

The following text is an excerpt from an email exchange with Robert Lordan, author of the Book ‘The Knowledge: Train Your brain Like a Cabbie’ (2018), who has given consent to be quoted on this:

> *“In terms of planning a route: When a passenger asked me a destination, usually the first thing that would happen is my brain would latch on to the compass point. So for example, say I picked up at St Pancras and somebody wanted to go to Brixton, my mind would immediately orientate its way south*.

> *How I’d then plan the route could depend. If it was a common destination-say for example Harrods or Waterloo station, I wouldn’t even have to think; my brain would be on autopilot. With big points it’s easy, like a moth drawn to a light!*

> *For longer, tricker routes, I’d find that my brain would often plan in stages; essentially I’d envision a set of waypoints and the route would then come to me as I progressed*.

> *In terms of how my general feeling towards London was impacted: The Knowledge made me crave detail! To this day I want to know as much as I can about London; what story and history lies behind every street*.

> *The city buzzes inside my head which is why I love to write about it. I already loved the city, but in studying it I now love it all the more. It feels like an old, familiar friend*.

> *The Knowledge also makes you want to know as much as you can about new locations that you’ve never been to before; if I go on holiday for example the first thing I do is study maps of the area as I want to know it and be able to get around said location as efficiently as possible once I’m there!”*

